# A method to build extended sequence context models of point mutations and indels

**DOI:** 10.1101/2021.12.06.471476

**Authors:** Jörn Bethune, April Kleppe, Søren Besenbacher

## Abstract

The mutation rate of a specific position in the human genome depends on the sequence context surrounding it. Modeling the mutation rate by estimating a rate for each possible k-mer, however, only works for small values of k since the data becomes too sparse for larger values of k. Here we propose a new method that solves this problem by grouping similar k-mers using IUPAC patterns. We refer to the method as k-mer pattern partition and have implemented it in a software package called kmerPaPa. We use a large set of human *de novo* mutations to show that this new method leads to improved prediction of mutation rates and makes it possible to create models using wider sequence contexts than previous studies. Revealing that for some mutation types, the mutation rate of a position is significantly affected by nucleotides that are up to four base pairs away. As the first method of its kind, it does not only predict rates for point mutations but also indels. We have additionally created a software package called Genovo that, given a k-mer pattern partition model, predicts the expected number of synonymous, missense, and other functional mutation types for each gene. Using this software, we show that the created mutation rate models increase the statistical power to detect genes containing disease-causing variants and to identify genes under strong constraint, e.g. haploinsufficient genes.

## Introduction

The germline mutation process is the source of all genetic variation, including all adaptive and deleterious variants. Understanding and modeling this process can be used to calibrate variant calling^1^, infer demographic history^2^, infer patterns of genome evolution^3^, identify sequences of clinical relevance for human diseases^4^, and to infer selective constraints of genes^5^. The main factor determining the mutation rate of a given site in the human genome is the sequence context surrounding the position. For example, spontaneous deamination of methylated cytosines results in ten times higher C-to-T rates at CpG sites^6–9^. Other factors that affect the mutation rate - such as GC content, CpG islands, epigenetic modifications - are associated with varied mutation rates depending on nucleotide context^10^ and thus cannot be studied without taking sequence context into account.

In this article, we introduce a new method to estimate mutation rates based on nucleotide context. Previous studies estimated such rates by assigning an independent rate to each k-mer^10,11^ or using a logistic regression model with a dummy variable for each nucleotide at each k-mer position^12^ and allowing interactions between at most four positions. Both these strategies have been used to build models that predict mutation rates using 7-mer contexts but become infeasible for longer contexts. Furthermore, these previous studies focused on point mutations, and little effort has been given to estimating the position-specific rate of germline indels. Neither Carlson et al.^10^ nor Aggarwala and Voight^12^ consider indels.

Samocha et al. calculate the rate of frame-shift indels per gene but do not model the indel mutation probabilities at each site^11^. Instead, they estimate the rate of frame-shift indels in a given gene by assuming that the rate of such variants is proportional to the rate of nonsense mutations. While some correlation exists between the number of polymorphic nonsense mutations and the number of polymorphic frame-shift indels, this correlation is primarily due to selection and not mutation, which makes the effectiveness of that strategy doubtful. Since frame-shift indels account for 44% of the LoF mutations in human genes^5^, making models that can predict the rates of such mutations should be considered an important task.

## Results

To overcome the problem that many k-mers will have few or zero observations if we use long k-mers to predict the mutation rates, we propose a new method that we call *k-mer Pattern Partition* (kmerPaPa). The main idea is that we partition the set of all k-mers by a set of IUPAC patterns so that each k-mer is matched by one and only one of the IUPAC patterns in the set. This partition should be done so that a pattern matches k-mers with similar mutation probabilities. Fig. 1 shows an example of how the 16 possible 3-mers containing C→G mutations can be partitioned using 10 patterns. An exact algorithm to calculate the pattern partition that optimizes the loss function is presented in the methods section. The loss function contains two regularizing hyperparameters *c* (complexity penalty), and *α* (pseudocount), which are fitted using 2-fold cross-validation.

**Fig. 1.**
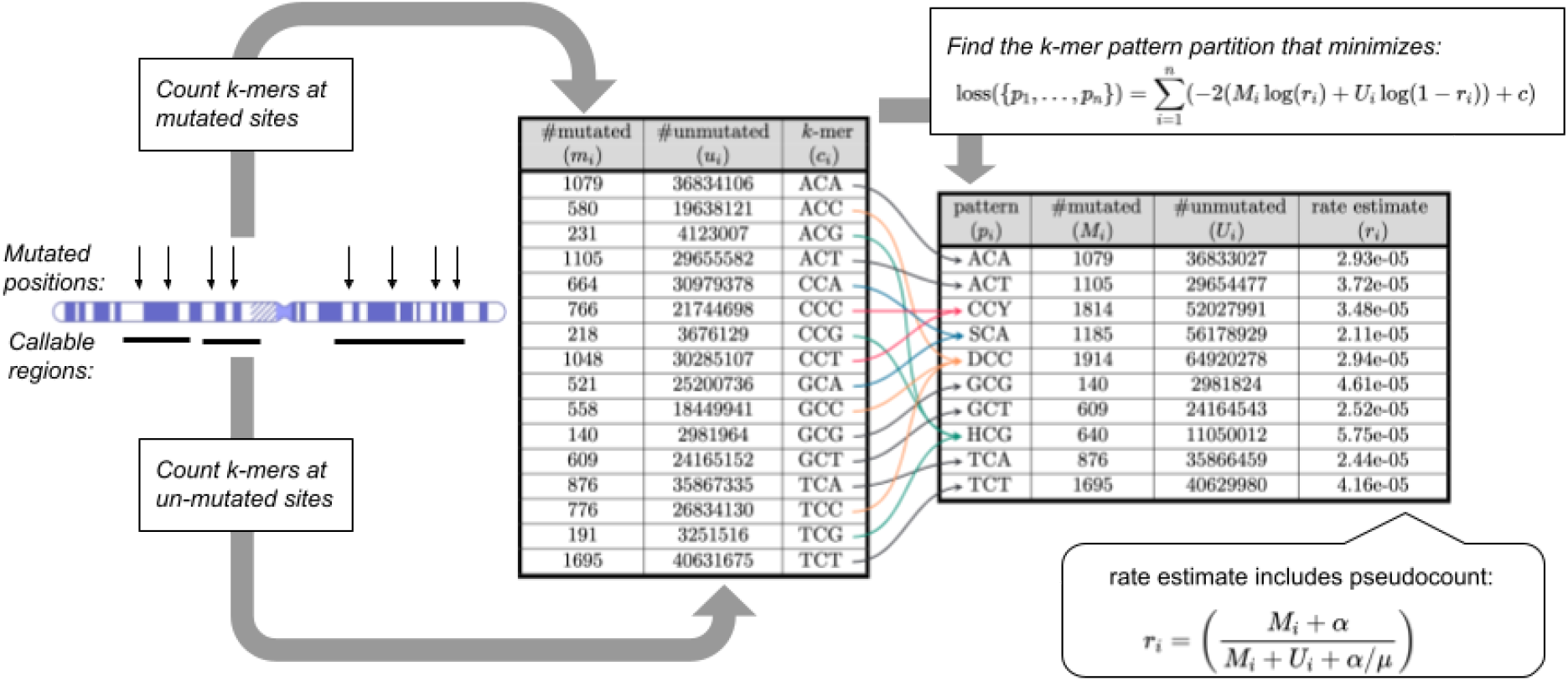
Overview of k-mer pattern partitioning (kmerPaPa) model. Input data consists of a list of observed mutations (here C→G mutations) and a bed file with regions sufficiently covered by WGS data to detect mutations. From this, we calculate a table with the number of times each possible k-mer is observed with the central base mutated and unmutated. The k-mers are then grouped using a set of IUPAC patterns so that each k-mer is matched by one and only one pattern. Out of the exponentially many possible pattern partitions the one that minimizes the loss function is chosen.

### Testing prediction of point mutation probabilities

To make models that only reflect the mutation rate and have as little bias from selection and biased gene conversion as possible, we use observed *de novo* mutations as training input to our model (see methods). To test the model, we first separate the data into a test and training set, using the even-numbered chromosomes for training and the odd-numbered as test data (see supplementary Fig. S1 for an overview of the data used in different analyses). We then fit independent pattern partition models to each of the six mutation types (A→C, A→G, A→T, C→A, C→G, C→T) for different values of k. We compared the performance of kmerPaPa to a model that assigns a rate to each k-mer (called “all k-mers” in Figure 2). As with kmerPaPa we also include a pseudocount in the “all k-mers” model and fit its value using 2-fold cross-validation. Figure 2a shows the test out-of-sample performance of these models for different values of k. For kmerPaPa, the joint Nagelkerke r^2 across the 6 mutation types keeps increasing as k increases. Whereas the alternative “all k-mers’’ model begins overfitting at 5-mers and performs poorly for larger values of k. The results reveal that nucleotides situated more than three base pairs away can affect a position’s mutation rate as the 9-mer partition outperforms the 7-mer partition in four of the six different mutation types, even though the 9-mer models tend to include fewer patterns (see supplementary Fig. S2).

**Fig. 2.**
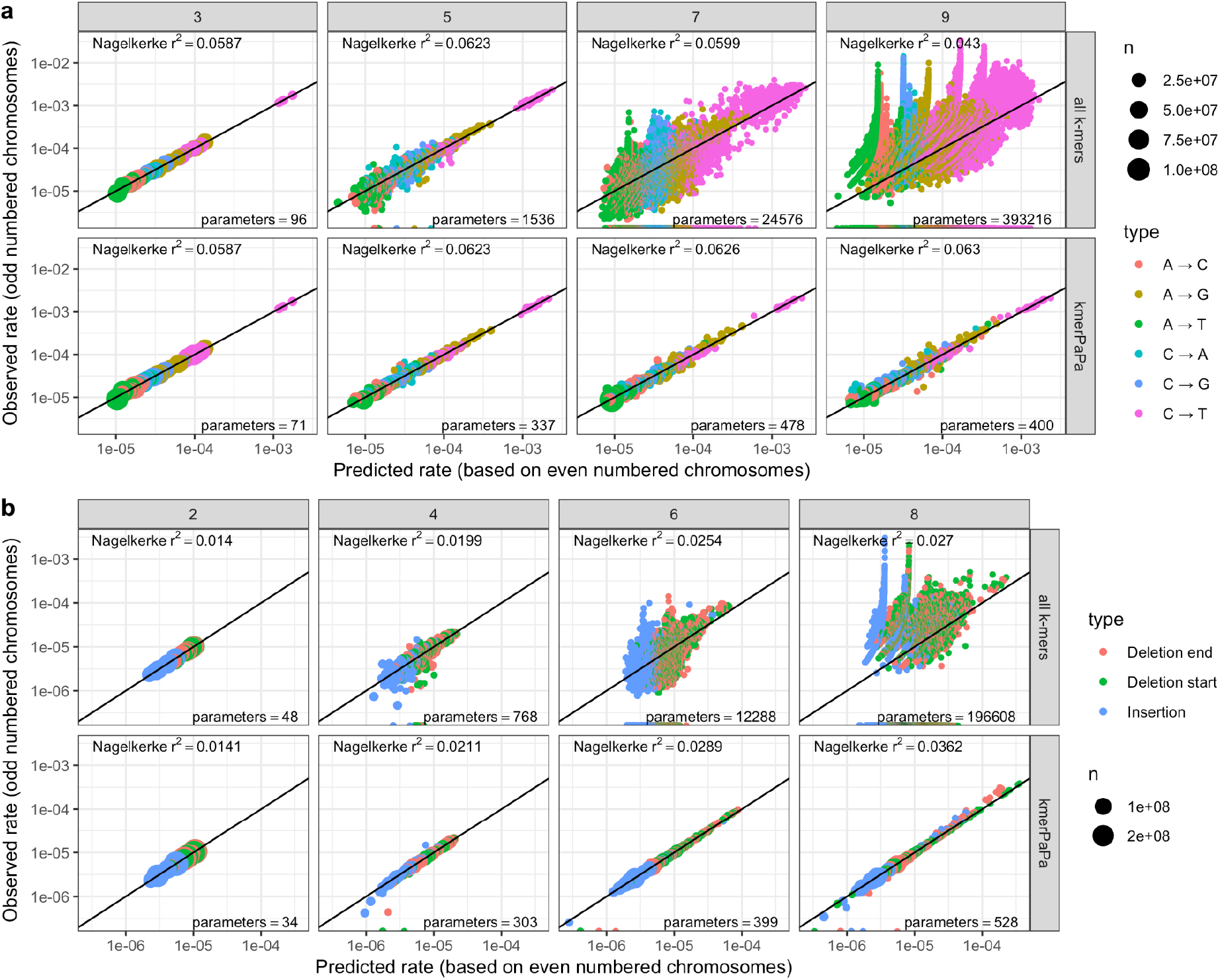
kmerPaPa performance on test data compared to predicting a rate per k-mer. All plots show predicted rates on training data (even-numbered chromosomes) on the x-axis and observed rates on test data (odd-numbered chromosomes) on y-axis. First row (“all k-mers”) show the rates for each k-mer. Second row (“kmerPaPa”) show the rate for each k-mer pattern. The sizes of the points reflect the number of times the k-mer/pattern is observed in the genome **a**) The predictions for point mutations for four different values of k. **b**) The predictions for indels for 4 different values of k.

### Testing prediction of indel mutation probabilities

Besides point mutations, it is essential to consider insertions and deletions. We assign mutation probabilities to indels by looking at the k-mers around the breakpoints - one for insertions and two for deletion (start and end). However, unlike point mutations, it is often impossible to precisely determine the position of an indel. Usually, more than one possible mutation event could have changed the reference sequence to the alternative sequence. If, for instance, “CAG” is changed to “CAAG,” it is impossible to know whether the DNA break and base insertion happened between C and A or between A and G. We handle this uncertainty by enumerating all the possible positions and randomly selecting one of them for each observed event (Figure 3). We evaluate the performance of the predicted indel rates using the same test/train split as we used for point mutations (Figure 2b). Both for all-kmers and kmerPaPa the out-of-sample performance increases with increased k. But the kmerPaPa model outperforms the all-kmers model for all values of k.

**Fig. 3.**
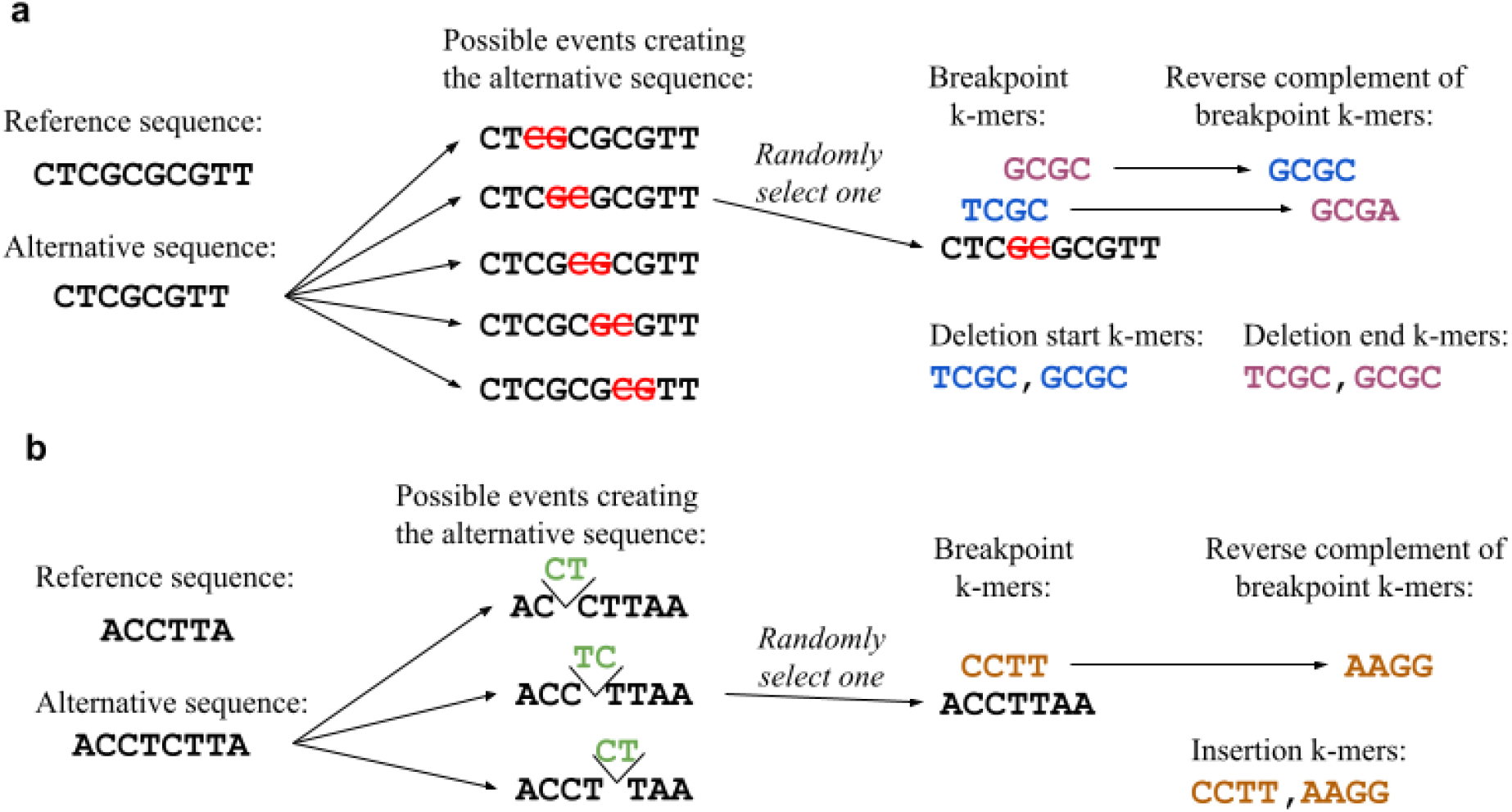
Counting k-mers for indels. The exact positions of indels are often indeterminable based on the sequence data. To handle this we enumerate all possible events creating the alternative sequence and select one of them at random. We do this independently for each indel mutation. Unlike point mutations, we count both the observed k-mer and its reverse complement. **a**) Deletion example. For deletions where we consider both the k-mer around the start breakpoint and the end breakpoint. The reverse complement of a deletion start k-mer is a deletion end k-mer and vice versa. **b**) Insertion example.

After validating that the extended k-mer models built by kmerPaPa give good results on an independent test set, we trained new kmerPaPa models on the whole data set. The following sections use these models to investigate the sequence contexts that increase mutation rates, find genes harboring disease-causing variants, and quantify the intolerance to mutations of human genes.

### Identification of patterns with unusually high or low mutation rates

An advantage of a k-mer pattern partition compared to a regression model is the direct interpretability of the output. By ordering the output patterns based on their mutation rate, we can directly observe patterns with extreme mutation rates. Figure 4 shows the relative mutation rate of each of the patterns for each of the mutation types (blue points) and the rate of each 3-mer (red points). For most mutation types the range of the predicted mutation types using k-mer patterns is an order of magnitude larger than those predicted by 3-mers. Many of the patterns with high point mutation probabilities match the patterns previously reported by Carlson et al.^10^ and Aggarwala and Voight^12^. For instance, the top pattern for A→G mutations “YYCA<u>A</u>TG” match the CCA<u>A</u>T motif reported by Carlson et al.^10^ and the YCA<u>A</u>TB pattern reported by Aggarwala and Voight^12^.

**Fig. 4.**
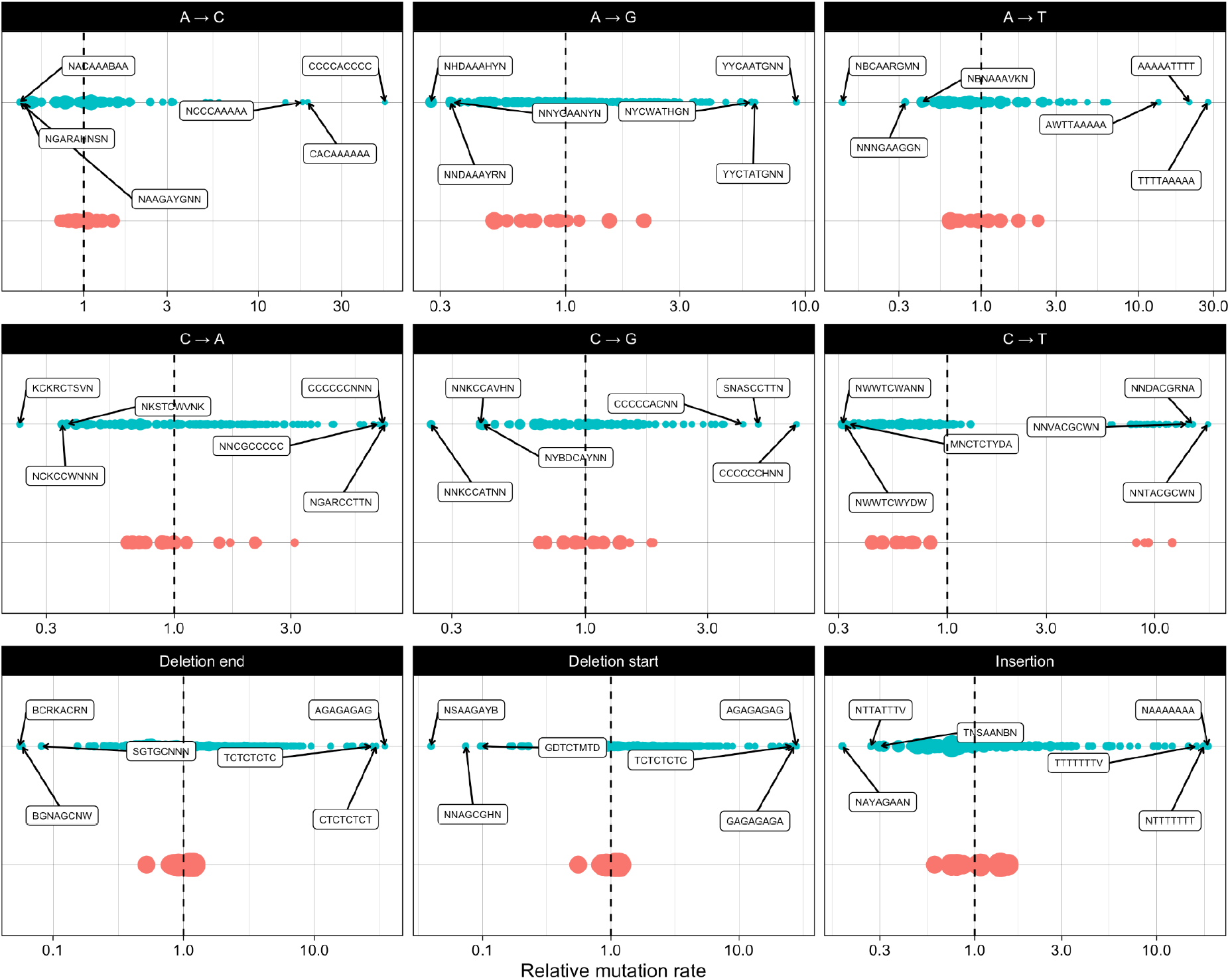
The relative mutation rate for each of the patterns in the kmerPaPa models. The blue points show the rate for the kmerPaPa patterns, the red points show the rate for each 3-mer. The dashed vertical line corresponds to the average mutation rate for the mutation type and the mutation rate values on the x-axis are relative to this average and on a log-scale. The size of the points reflect the number of genomic sites that match the pattern.

Previous studies have reported that polymerase slippage at short tandem repeats is responsible for 75% of indels^13^. It is thus not a surprise that many of the indel patterns we observe contain repeated sequences. For insertions, we observe the highest mutation rate for (T)n and (A)n mononucleotide repeats followed by (G)n and (C)n repeats. For the deletions, the top patterns correspond to (AG)n and (CT)n di-nucleotide repeats. After these, we observe a high deletion rate of the middle two (underlined) bases in the palindromic sequence: CAC<u>AT</u>GTG. This sequence has not previously been described as a mutation hotspot and the reason behind its high deletion rate is unknown. One possible explanation is that the palindromic sequence can form a hairpin structure. DNA hairpins are known to cause transient polymerase stalling, which leads to indel formation, whereas other alternative DNA structures (e.g. G4) cause persistent polymerase stalling and result in point mutations^14^.

### Detection of genes where *de novo* mutations cause disease

Genes in which germline mutations cause disease can be found by looking for genes with surprisingly many *de novo* mutations in afflicted children. The k-mer pattern partitions we have created can be used to calculate how many mutations of a specific functional category (synonymous, missense, etc.) to expect in a given gene.

We have created a software tool - Genovo - to enumerate all the possible variants of a specific functional category and look up their mutation rates. Given a list of observed mutations, Genovo can then calculate p-values by sampling from the Poisson-Binomial distribution given by this list of rates. This functionality is similar to that provided by the tool denovolyzeR^15^, so we compare our results to this tool.

As a first test, we compare the genic mutation rate predicted by Genovo and denovolyzeR to the number of segregating variants in each gene. First, we look at the number of rare (MAF <1%) synonymous variants for each gene in the gnomAD database, where we observe a better correlation with the synonymous rate per gene estimated by Genovo (r=0.977) than by denovolyzeR (r=0.958) (Fig. 5a). Most of the variation in the number of synonymous variants per gene can be explained purely by the genes’ length. But if we look at the rate per site and divide both the observed number and the predicted rate by the length of the coding sequence, we still observe high correlations (r=0.699 for Genovo and r=0.679 for denovolyzeR) (Fig. 5b). For nonsynonymous variants, we expect smaller correlations since differences in selective pressure between genes greatly influence these. For all classes, we observe better correlations with the rates predicted by Genovo than those predicted by denovolyzeR.

**Fig 5.**
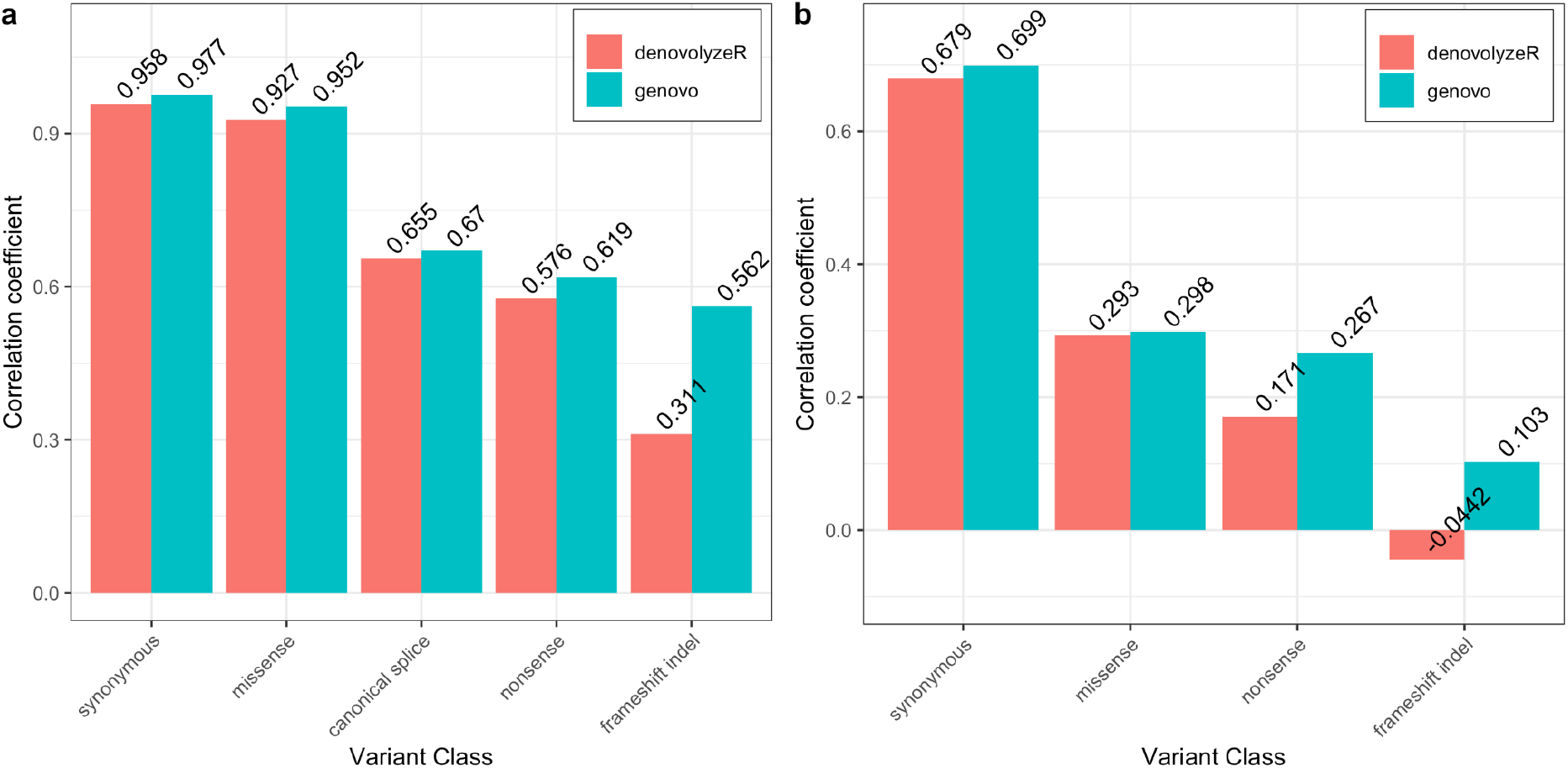
Correlation between the predicted genic rate for a mutation type and the number of segregating variants of that type. **a**) Pearson correlation coefficient between the observed number of mutations per gene and the expected number using Genovo and denovolyzeR. **b**) Pearson correlation coefficient between the observed number of mutations per coding position and the expected number per coding position for each gene using Genovo and denovolyzeR.

Secondly, we want to compare the statistical power of Genovo and denovolyzeR when it comes to identifying disease-causing genes. To do this, we look at 4293 trios from the Deciphering Developmental Disorders consortium^16^ and divide the set of trios into an equally sized test and train data set. 350 genes contain at least one loss-of-function(LoF) mutation in the test data set. 45 of these show significant LoF enrichment after Bonferroni correction according to Genovo, and 33 of these are significant in the test data set. Running denovolyzeR yields 34 significant genes in the train set, 24 of which are significant in the test set. This means that Genovo has a higher validation rate in the independent test data set (73.3% compared to 70.6%) even though it identified a larger number of significant genes in the training data set.

### Quantifying genic tolerance to Loss of Function mutations

The ability to predict the number of expected mutations in a gene is not only relevant for finding genes where a specific cohort has an enriched number of functional mutations. We can also use it to find genes that contain fewer segregating variants than expected because deleterious variants have been purged from the population by selection. The types of variants that are most likely to be deleterious are those that we expect to completely inactivate a gene, such as stop-gain, essential splice, and frameshift variants. The observed number of such loss-of-function (LoF) variants for a gene divided by the expected number predicted by our mutation rate model - the LoF O/E ratio - provides an estimate of evolutionary constraint. If a gene has a LoF O/E ratios around one it indicates that it is evolving neutrally and that no substantial fitness cost is associated with losing the gene. In contrast, haploinsufficient genes where two functional alleles are essential for survival will have LoF O/E ratios at or close to zero. We have calculated this ratio for each gene using the observed mutations from gnomADv2 and compared them to the ratios reported for each gene by gnomAD. Looking at genes predicted to be haploinsufficient by ClinGen^17^ we observe that 55.5 percent of the genes have a Genovo O/E ratio in the first decile compared to 47 percent for O/E ratio calculated by gnomAD (see Fig. 6a). If we instead of the LoF O/E ratio use the upper bound of the 90 confidence interval of the ratio (LOEUF) as suggested by Karczewski et al^5^ we see similar results (see Fig. 6b). We also compared Genovo and gnomAD using two lists of essential genes used in the gnomAD article^5^ and obtained similar results (see supplementary Fig. S4). If we look at the ability to classify whether a gene is haploinsufficient we get significantly higher AUC values using Genovo (p-value: 2.42 e-153, See Fig. 6c).

**Fig 6.**
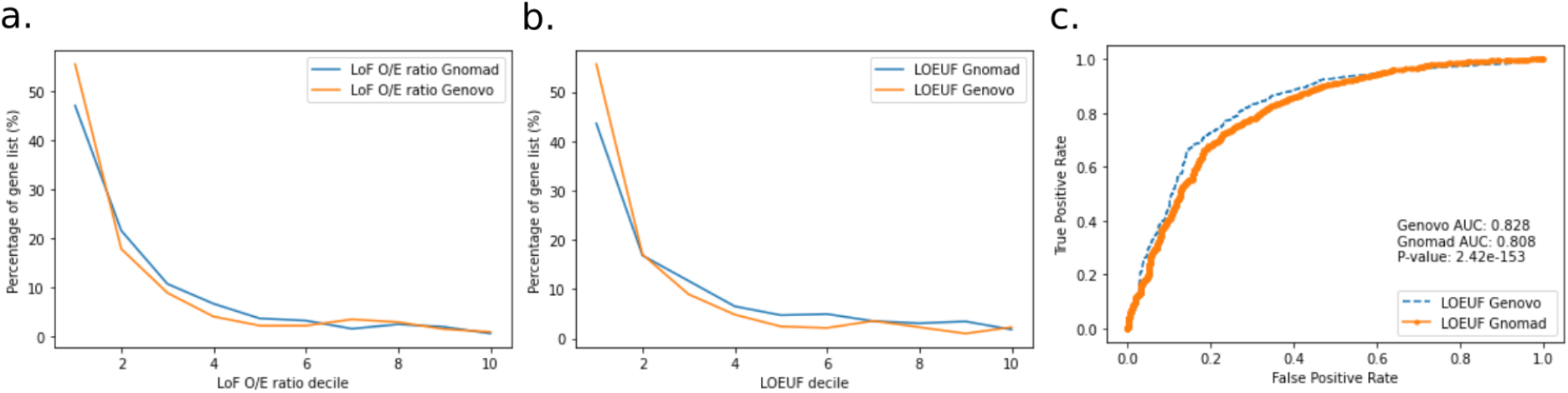
Inferring constraint of haploinsufficient genes. **a**) Comparison of observed/expected ratio for loss of function variants (LoF O/E ratio) inferred by Genovo and gnomAD (blue and orange, respectively). X-axis depicts LoF O/E ratios sorted into deciles, whereas the y-axis depicts the percentage of genes in each decile. **b**) Comparison of categorisation by LOEUF scores inferred by Genovo - and LOEUF score by gnomAD. X-axis depict LOEUF scores sorted into deciles, whereas the y-axis depicts the percentage of genes in each decile. **c**) Receiver Operating Characteristic (ROC) curve comparing the ability to predict whether a gene is annotated as haploinsufficient using LOEUF scores from gnomAD and Genovo.

## Discussion

The k-mer pattern partition method introduced in this manuscript makes it possible to build robust models of the germline mutation process using k-mers. We have demonstrated the method by building models for each of the six point mutation types as well as short insertions and deletions using *de novo* mutations as input. The results show that the out-of-sample error decreases as k increases proving that even nucleotides four bp away from a given position are informative about its mutation probability. An advantage of the pattern partition models is their interpretability, with overrepresented patterns readily available not requiring any further analysis of the created models. The k-mer pattern partition approach is general and could, in principle, also be used to solve other problems where the input is small position-specific k-mers. The main shortcoming of the method in its current form is that the presented dynamic programming algorithm is exponential in running time which makes it infeasible to use for k-mers longer than nine bases. A possible improvement to alleviate this problem would be to consider heuristic algorithms that could quickly find a good (but not necessarily optimal) solution and thus enable analyses with larger k’s.

The predictive models created using kmerPaPa are not only robust, they have the added benefit that they are easily interpretable. The method can be seen as a dimensionality reduction method that reduces the high dimensional k-mer space to a lower dimension space where each dimension is represented by a IUPAC pattern. This set of patterns with associated mutation rates makes it very easy to see the general differences between the high mutation rate k-mers and low mutation rate k-mers. Other methods have previously been used to create models of point mutation rates based on sequence context but it is novel that we in this study also create robust models of indel mutation rates based on sequence context. To do that, we had to overcome the problem that the exact position of an indel often is indeterminable. We do this by enumerating all the possible positions and randomly choosing one of them for each indel. Future studies that train indel kmerPaPa models using larger input data sets and further examine the resulting patterns can hopefully increase our understanding of the indel mutation process.

Besides the k-mer pattern method implemented in the kmerPaPa software, we also present a tool called Genovo that can calculate the expected number of mutations with a specific functional consequence in a gene. Furthermore, Genovo can also calculate p-values for whether a gene contains significantly more observed mutations of a particular functional category than expected. This functionality can be used to find genes with causative *de novo* mutations in disorders such as autism and developmental disorders. We have compared Genovo to denovolyzeR, which contains similar functionality. Trying to predict the number of synonymous mutations in each gene reported in gnomAD we see that we do slightly better than denovolyzeR. Our ability to predict the number of frameshift mutations in a gene is, however, much better than denovolyzeR. This reflects that we train independent mutation rate models for insertions and deletions whereas denovolyzeR calculates the expected number of indel variants by assuming a correlation with the number of point mutations. Unlike denovolyzeR, which is based on a specific reference genome (hg19) and gene annotation, Genovo is more general and can calculate expectations and p-values for any reference genome and gene annotation. This allows users to use the newest reference genome and gene annotation at any time and makes it possible to use kmerPaPa and Genovo to analyze enrichment or depletion of functional mutational categories in other species.

In conclusion, we have created a new robust method for predicting mutation probabilities based on sequence context for both point mutations and indels. Furthermore, we have developed a flexible software tool that makes it easy to use the created mutation rate models to find genes where *de novo* mutations cause disease or to measure the evolutionary constraint on genes.

## Methods

### Definition of k-mer pattern partition

A k-mer pattern partition is a set of IUPAC-encoded nucleotide patterns that partitions a set of k-mers so that each k-mer is matched by one and only one of the IUPAC patterns in the set. To find a pattern partition that groups k-mers with similar mutation rates as well as possible while not overfitting the data, we select the partition P= {p_1_, …, p_l_} that optimizes the following loss function:

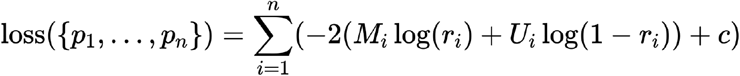

Where M_i_ and U_i_ are the number of mutated and unmutated positions that match the i’th pattern (p_i_), and c is a regularizing complexity penalty. And r_i_ is the rate estimate for sites that match the i’th pattern regularized to be closer to the mean mutation rate, *μ*, using a pseudo count, *α*:

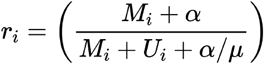

The two regularizing hyperparameters c, and *α* and fitted using cross-validation.

### Calculating the optimal pattern partition

Starting with the most general pattern, we can create any possible pattern partition by:

1. Picking a position in the pattern.
2. Splitting the IUPAC code at that position into two subcodes that form a partition (we call such a split a two-partition).
3. Possibly repeating these steps on each of the two sub-patterns.

Table 1 shows all the possible two-partitions for each IUPAC code.

**Table 1.** The definition of all IUPAC characters and the list of all possible two-partitions for each IUPAC character.

| IUPAC code | Set | Two-partitions |
| --- | --- | --- |
| S | {C,G} | (C,G) |
| W | {A,T} | (A,T) |
| R | {A,G} | (A,G) |
| Y | {C,T} | (C,T) |
| K | {G,T} | (G,T) |
| M | {A,C} | (A,C) |
| B | {C,G,T} | (C,G), (G,Y), (T,S) |
| D | {A,G,T} | (A,K), (G,W), (T,R) |
| H | {A,C,T} | (A,Y), (C,W), (T,M) |
| V | {A,C,G} | (A,S), (C,R), (G,M) |
| N | {A,C,G,T} | (A,B), (C,D), (G,H), (T,V), (R,Y), (S,W), (K,M) |

Since all pattern partitions can be created using the strategy above, we can find the minimal loss for the pattern, p, using the following recursion formula:

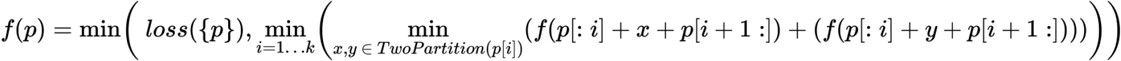

Using this formula, we compute the optimal partitions bottom-up using dynamic programming so that the optimal partition for a given pattern is never calculated more than once. This algorithm’s running time and memory usage are proportional to the number of possible patterns - and thus exponential in k. Our python implementation, speeded up using numba^18^ for just-in-time compilation, can calculate the optimal 9-mer pattern partition for C→T mutations in ∼4 hours (given *α* and *c*). If we wanted to calculate the optimal 11-mer, we would, however, need to multiply this with 15^2 = 225, so that is not feasible using this algorithm. To find the optimal *α* and *c* we do cross-validation over a grid and this means that we need to calculate the optimal pattern for each fold and each *α* and *c* combination. This grid search is, however, easy to parallelize since each parameter combination can be run separately on different machines.

### De Novo Mutations and covered regions

To train kmerPaPa models, we use *de novo* mutations from 6 different WGS studies of parent-offspring trios^19–24^. Together these studies include 379330 autosomal point mutations from 7206 trios. Four of the six studies also report indel mutations (39877 autosomal variants). Studies mapped to hg19 were first lifted to hg38.

Detection of *de novo* mutations requires strict filters and usually, there will be a substantial fraction of the genome where *de novo* mutations cannot be called due to insufficient coverage or low complexity sequence. We do not know the exact genomic regions where a mutation could have been called in each of the six studies. To deal with this, we have chosen a conservative set of regions - the strict callable mask from the 1000 genomes project^25^ - and assume that mutations could have been called in this subset of the genome in all of the studies. Because we want to infer a genome-wide mutation model, we further exclude the C→G enriched regions described by deCODE^26^ since these are known to have mutation patterns that differ substantially from the average. After removing the C→G enriched regions from the 1000g callable, the callable regions contain 1754 million autosomal sites and after discarding all mutations that fall outside these regions, we are left with 249437 autosomal SNVs and 22331 autosomal indels.

We use the program kmer_counter (https://github.com/BesenbacherLab/kmer_counter) to calculate the k-mer counts used as input kmerPaPa. As explained in the results section we first test the kmerPaPa method by using the even-numbered chromosomes as training data and the odd-numbered chromosomes as test data. Afterwards, we train models using mutations on all chromosomes (see Supplementary Fig. S1). Besides training indel models using all insertions and all deletions as presented in Fig. 4 we also train separate models for indels that are a multiple of 3 and indels that are not a multiple of 3. These models are used as input for the Genovo software to allow it to calculate separate probabilities for frame-shift and in-frame indels.

### Genovo

From a list of observed mutations the Genovo software can determine the number of 1) synonymous, 2) missense, 3) nonsense, 4) start-codon disruption, 5) stop-codon disruption, 6) canonical splice site disruption 7) inframe indel and 8) frame-shift indel events in each transcript. Furthermore, the software takes the genomic sequence of every transcript and enumerates all possible point mutations. For each possible point mutation, it determines the type of point mutation and its probability derived from our kmerPaPa models. The possible mutations of each mutation type are tallied up to calculate the expected number of point mutations of each mutation type. In addition, the possible mutations and their probabilities are used to calculate a Poisson-binomial distribution of mutation events for every transcript, because each possible mutation event has a different probability to occur. We then sample from this distribution to calculate the p-value for the number of observed mutations of each mutation type and to calculate the confidence interval around the number of expected mutations (see Supplementary Fig. S3.)

### Scaling mutation rates

The mutation rates output by kmer-PaPa corresponds to the number of mutations expected in the number of individuals that we have data from. To turn these estimates into rates per generation we multiply all SNV rates with a factor so that the average mutation rate becomes 1.28×10^×8 26^. And we scale indel rates to have an average rate per site of 0.68×10^×9 27^. In Genovo these mutation rates should then be scaled to reflect the number of generations that the observed mutations correspond to. To avoid that some mutations get probabilities above one for data sets with many samples we use the following formula to scale the rate per base per generation, *r*, to get the probability that this mutation has occurred in n_gen_ observed generations:

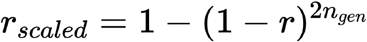

For the gnomAD analysis where we do not have *de novo* mutations from a certain number of trios but segregating variants accumulated over an unknown number of generations, we set *n*_*gen*_ so that the number of expected synonymous mutations fit the observed number of synonymous mutations in the whole exome.

### Correction for coverage

Coverage bias in exome sequencing data results in fewer sequenced individuals for some transcripts compared to others in the gnomADv2 data. This means that the expected number of mutations predicted by Genovo will be too high for genes that have low coverage. To correct for this we fit a lowess curve using the statsmodels package ^28^ in python, to a scatter plot with synonymous observed/expected values (y-axis) vs. the number of called samples (x-values) for each transcript (see Supplementary Fig. S5). The number of called samples for a transcript was estimated as the average AN (from the gnomAD2 vcf file) over all variants in the transcript. In the lowess-function, we used 0.05 as the fraction of the data used when estimating each y-value. Based on this inference, we made a cut-off where we only kept transcripts that have an average AN value above 50000. For each transcript, we then divided the expected variant count with the corresponding lowess estimate for all variant classes to get a coverage corrected expectation.

### Running Genovo

To produce the expected variant count for each transcript we ran Genovo using hg19 and gencode v. 19 (same as used for gnomADv2) using a scaling factor of 2.2×10^−7^. Then we applied the coverage correction as described above. For the comparison with denovolyzeR shown in fig. 5 we used the VEP annotations provided in “gnomad.exomes.r2.1.1.sites.vcf” to get the observed number of variants in each functional class.

Of the 4293 trios in the DDD data set 3664 contain at least one exome mutation. When splitting the data into test and training data sets we put half of the 3664 individuals with a mutation in each group. When calculating p-values using Genovo we did it using 10 million random samples from the null model. For both Genovo and denovolyzeR we do bonferroni correction of the training set p-values based on the number of genes that have at least one observed LoF mutation in the training data (350 genes). For each gene we only test the transcript used by denovolyzeR when running Genovo.

### Calculation of Genovo LOEUF score

When calculating the “loss-of-function observed/expected upper bound fraction” (LOEUF score) ^5^ from the output of the Genovo software we do it by dividing the number observed LoF variants by the lower bound of the 90% confidence interval of the expected number of LoF variants. Like the other expected values this lower bound used in the denominator had also been adjusted for coverage differences between transcripts as described above in the section *Correction for coverage*. As there were many transcripts with zero observed mutations, we furthermore added a pseudo score of 0.5 to all values of observed and expected LoF.

### gnomAD and Genovo comparison

We used the pre-calculated LoF O/E ratios and LOEUF scores provided in the publicly available data provided by the gnomAD consortium (gnomad.v2.1.1.lof_metrics.by_transcript.txt.bgz) to compare O/E ratios and LOEUF scores between gnomAD and Genovo. We made sure that we compared transcripts that we had data for in both gnomAD and Genovo datasets. Thereafter, we split the transcripts into deciles and inferred how many transcripts were found in each decile. We did this for each model - gnomAD or Genovo - and visually compared the two for three different groups of genes that were: essentiality of genes inferred by two lists (mouse gene knockout experiments and cellular inactivation assays) and haploinsufficient genes. We used the same curated gene lists that were used in used in the gnomAD paper^5^ downloaded from https://github.com/macarthur-lab/gene_lists; haploinsufficient genes according to ClinGen Dosage Sensitivity Map (as of Sep 13 2018)^17^, genes deemed essential in mouse^29–31^, genes deemed essential in human^32^, and genes deemed essential and non-essential in human by CRISPR screening^33^.

We conducted the ROC AUC analysis by using the sklearn.metrics.roc_curve function from the scikit-learn module^34^ in python. For each of the AUC scores yielded for each prediction score (gnomAD or Genovo), we also conducted a DeLong test^35^ to infer whether the AUC scores were significantly different. To conduct this test we used python code adapted from https://github.com/Netflix/vmaf/.

## Data Availability

All input *de novo* mutation data sets used as input are publicly available. Two of them can be found as supplementary tables in the articles reporting them^19,20^. Two are available using links to external files provided in the articles^21,24^. The last two data sets^22,23^ can be downloaded from denovoDB^36^.

The kmerPaPa models trained on even-numbered chromosomes are available in Supplementary table 4. The kmerPaPa models trained on all chromosomes are available in Supplementary table 5. These models formatted as input files to Genovo can furthermore be found at https://github.com/BesenbacherLab/Genovo_Input. The predicted number of variants in each functional type for each gencode v19 transcript used for the gnomAD comparison are in Supplementary. table 6.

## Code Availability

kmerPaPa can be installed from https://pypi.org/project/kmerpapa/ and the source code is available at https://github.com/BesenbacherLab/kmerPaPa.

Genovo can be installed from https://crates.io/crates/genovo and the source code is available at https://github.com/BesenbacherLab/genovo.

kmer_counter can be installed from https://pypi.org/project/kmer-counter/ and the source code is available at https://github.com/BesenbacherLab/kmer_counter.

## Author Contributions

SB designed and implemented the kmerPaPa software. JB designed and implemented the Genovo software. SB ran the kmerPaPa software to produce mutation rate models and the Genovo software to produce predicted rates per mutation type per transcript. AK performed coverage correction and the gnomAD comparison analysis. SB and AK wrote the manuscript. All authors reviewed and approved the final manuscript.

## Acknowledgements

This work was funded by a Sapere Aude - Starting Grant (no. 4181-00490) from the Independent Research Fund Denmark.

## Supplementary Figures

**Supplementary Fig. S1.**
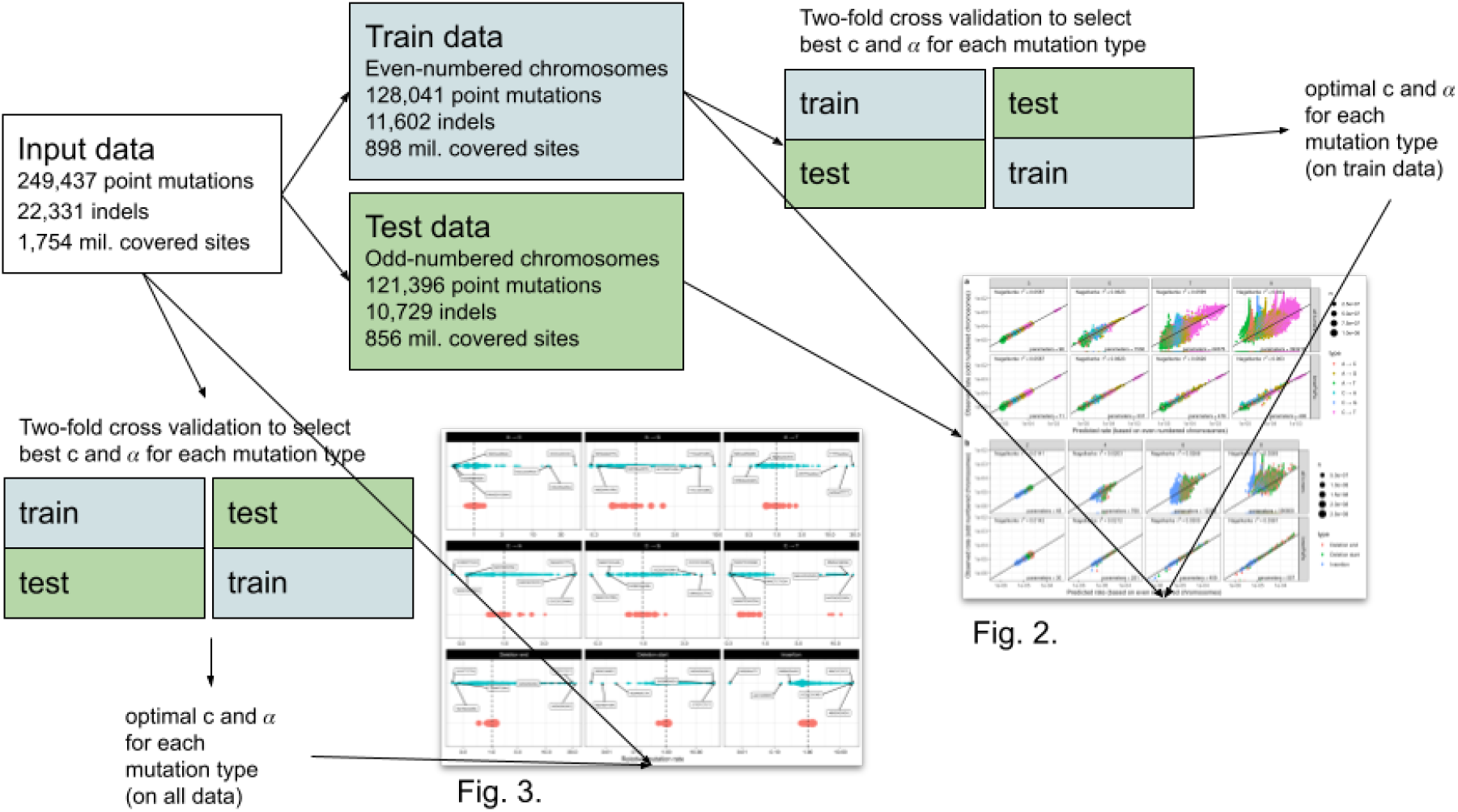
Overview of the data used in different analyses.

**Supplementary Fig. S2.**
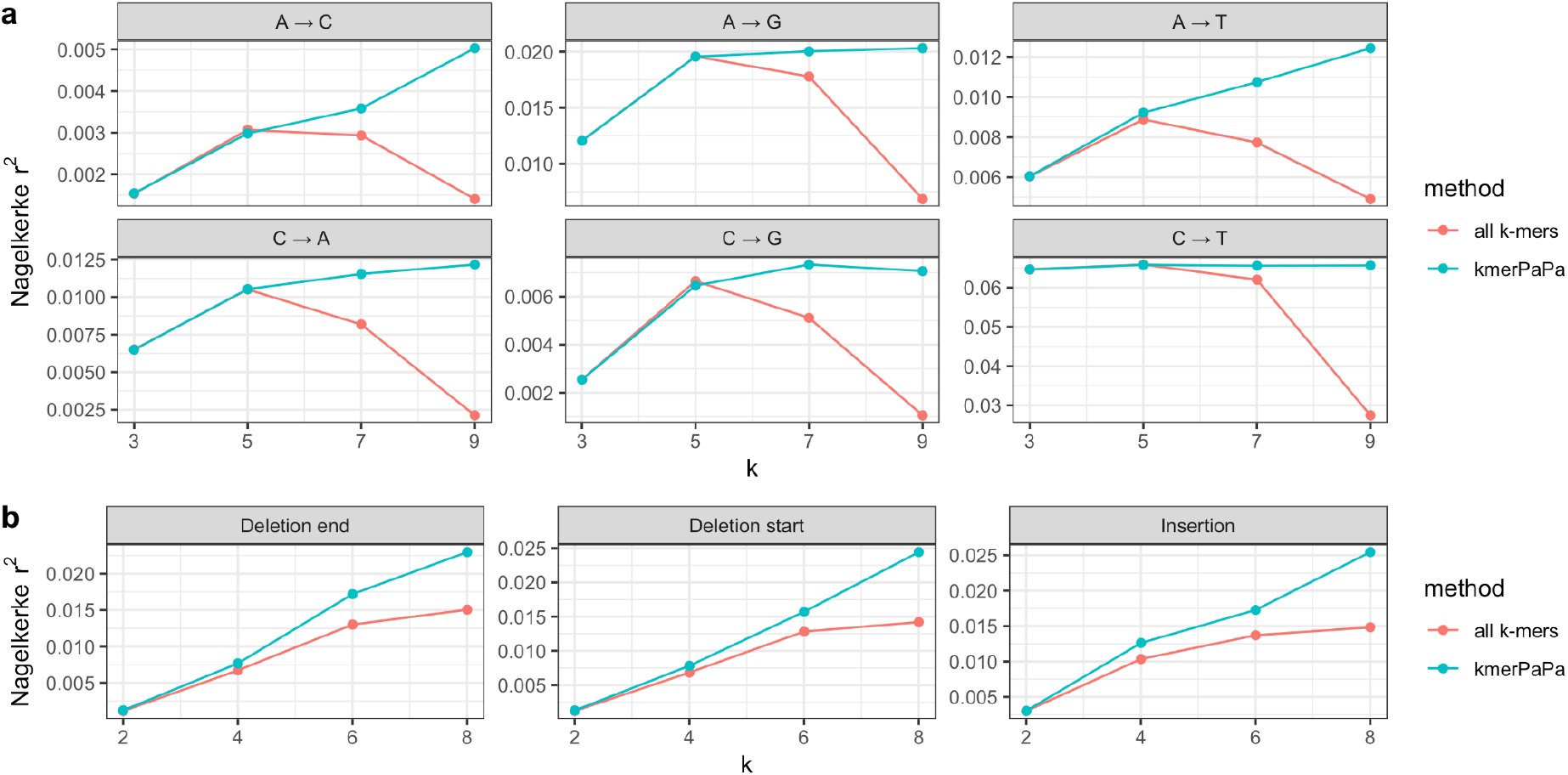
Nagelkerke r^2^ on test data for different values of k. For longer k-mers that “all k-mers” models overfit resulting in worse r^2^ values on the test data. This is not the case for the kmerPaPa models.

**Supplementary Fig. S3.**
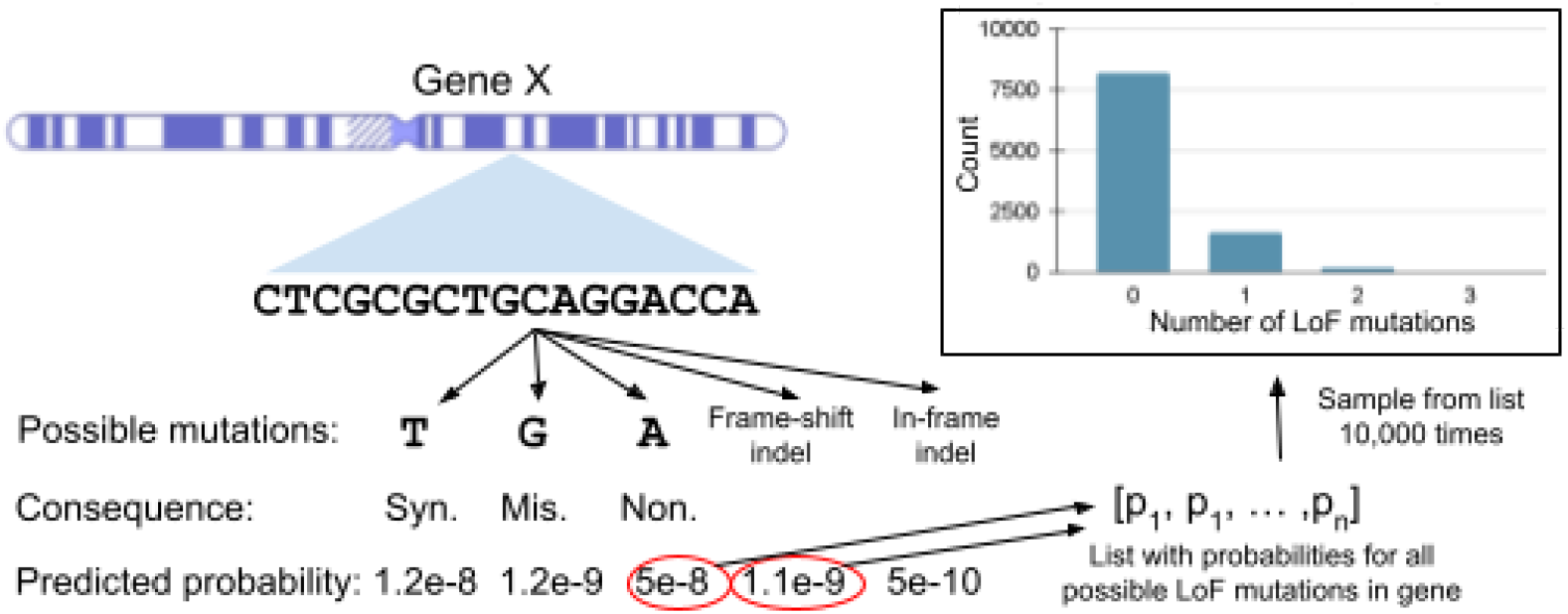
Calculating the null distribution for the number of expected LoF mutations in a gene. Genovo considers all possible mutations in a gene and calculates their functional consequence. For a given mutation type (fx. LoF) it will then calculate a list of the probabilities for all possible mutations of this type. Genovo then samples from this list to create the null distribution that is used to calculate the p-values.

**Supplementary Fig. S4.**
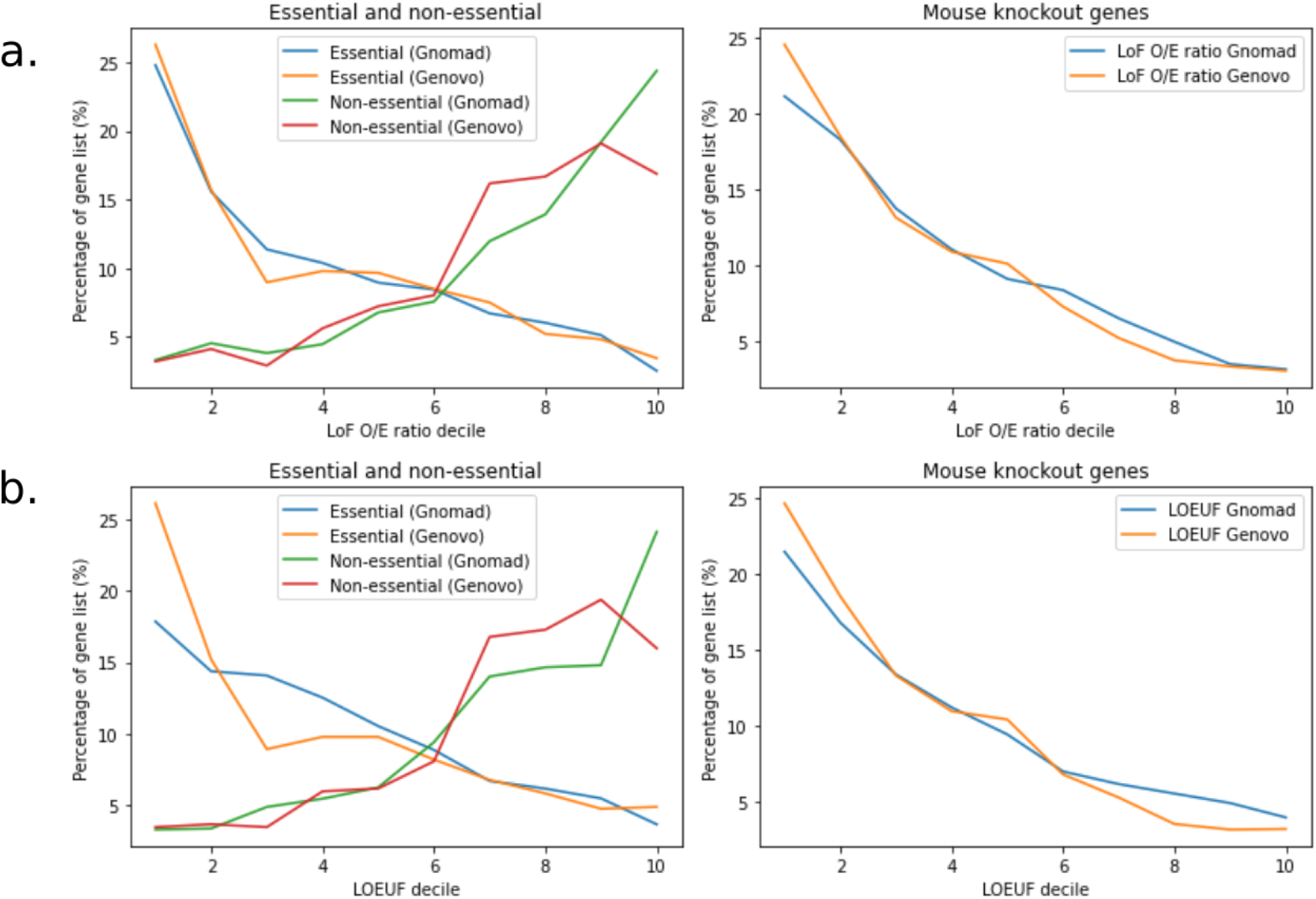
Results from LOEUF analysis. In the first column we observe essential and non-essential genes. The green and blue lines depict the percentage of essential genes for Genovo and gnomAD, respectively. The red and yellow lines display distribution of non-essential genes for Genovo and gnomAD, respectively. The second column depicts essential knock-out genes for mice, and the third column depicts haploinsufficient genes. Panel **a** displays LoF observed/expected ratio, panel **b** depicts LOEUF score as inferred by gnomAD and Genovo.

**Supplementary Fig. S5.**
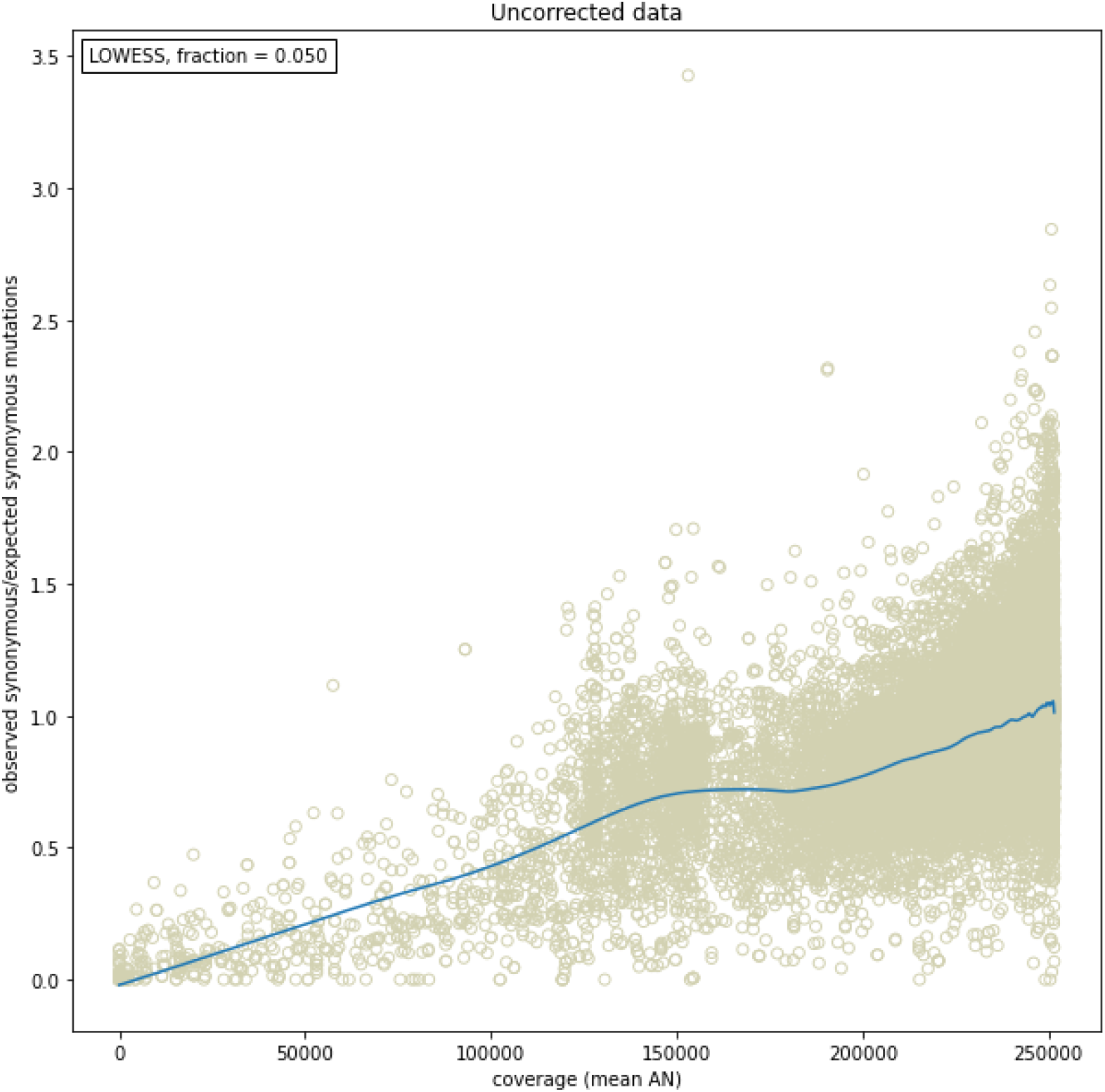
Coverage Correction. Smoothed lowess values (blue line) based from synonymous observed/expected mutation data, over coverage (beige data). ‘Fraction’ is a number between 0 and 1, which represents the fraction of the data used when estimating each y-value.

**Supplementary Table 1.** Data sets with *de novo* mutations used as input to kmerPaPa.

| Study | Trios | Autosomal point mutations | Autosomal indels | Reference Genome |
| --- | --- | --- | --- | --- |
| Halldorsson et al. 2019 | 2977 | 194687 | 13367 | hg38 |
| Goldmann et al. 2016 | 816 | 35748 | 0 | hg19 |
| Francioli et al. 2015 | 258 | 11016 | 0 | hg19 |
| Sasani et al. 2019 | 471 | 26794 | 1892 | hg19 |
| Yuen et al. 2017 | 1652 | 104948 | 16582 | hg19 |
| Turner et al. 2017 | 1032 | 109433 | 8036 | hg19 |

**Supplementary Table 2.** Percentage of haploinsufficient genes in each decile. Deciles clustered by Observed/Expected (O/E) ratio of Loss Of Function (LoF) mutations observed in haploinsufficient genes. Low numbered deciles (e.g. decile 1,2 or 3) contain the lowest O/E of LoF scores, whereas high numbered deciles contain higher O/E of LoF scores. These are the values plotted in Fig 6a.

| Decile | % gnomAD | % Genovo |
| --- | --- | --- |
| 1 | 47.0 | 55.5 |
| 2 | 21.6 | 17.9 |
| 3 | 10.8 | 8.9 |
| 4 | 6.7 | 4.1 |
| 5 | 3.7 | 2.3 |
| 6 | 3.3 | 2.3 |
| 7 | 1.6 | 3.6 |
| 8 | 2.5 | 2.9 |
| 9 | 2.0 | 1.6 |
| 10 | 0.7 | 0.9 |

**Supplementary Table 3.** Percentage of haploinsufficient genes in each LOEUF decile. Low numbered deciles (e.g. decile 1,2,3 …) contain genes with the lowest LOEUF scores, whereas high numbered deciles contain higher LOEUF scores. These are the values plotted in Fig 6b.

| Decile | % gnomAD | % Genovo |
| --- | --- | --- |
| 1 | 43.6 | 55.6 |
| 2 | 16.8 | 17.0 |
| 3 | 11.7 | 8.9 |
| 4 | 6.5 | 4.8 |
| 5 | 4.7 | 2.4 |
| 6 | 4.9 | 2.1 |
| 7 | 3.5 | 3.6 |
| 8 | 3.1 | 2.3 |
| 9 | 3.4 | 0.9 |
| 10 | 1.8 | 2.3 |

